# Jamestown Canyon Virus is transmissible by *Aedes aegypti* and is only moderately blocked by *Wolbachia* co-infection

**DOI:** 10.1101/2023.05.11.540455

**Authors:** Meng-Jia Lau, Heverton LC Dutra, Matthew J Jones, Brianna P McNulty, Anastacia M Diaz, Fhallon Ware-Gilmore, Elizabeth A McGraw

## Abstract

Jamestown Canyon Virus (JCV), a negative-sense arbovirus, is increasingly common in the upper Midwest of the USA. Transmitted by a range of mosquito genera, JCV has at its primary amplifying host, white-tailed deer. *Aedes aegypti* is the major transmitter globally of the positive-sense viruses dengue (DENV), Zika, chikungunya, and Yellow Fever. *Ae. aegypti’s* distribution, once confined to the tropics, is expanding, in part due to climate change. *Wolbachia*, an insect endosymbiont, limits the replication of co-infecting viruses inside insects. The release and spread of the symbiont into *Ae. aegypti* populations has been effective in reducing transmission of dengue and other viruses to humans. The mechanism of *Wolbachia*-mediated viral blocking in vectors is still poorly understood, however. Here we explored JCV infection potential in *Ae. aegypti*, the nature of the vector’s immune response, and interactions with *Wolbachia* infection. We show that *Ae. aegypti* is highly competent for JCV, growing to substantial loads and rapidly reaching the saliva after an infectious blood meal. The mosquito immune system responds with strong induction of RNAi and JAK/STAT. Neither the direct effect of viral infection nor the energetic investment in immunity appears to affect mosquito longevity. *Wolbachia* infection blocked JCV only in the early stages of infection. *Wolbachia*-induced immunity was small compared to that of JCV, suggesting innate immune priming does not likely explain blocking. We propose two models to explain why *Wolbachia’s* blocking of negative-sense viruses like JCV may be less than that of positive-sense viruses, relating to the slowdown of host protein synthesis and the triggering of interferon-like factors like Vago. In conclusion, we highlight the risk for increased human disease with the predicted future overlap of *Ae. aegypti* and JCV ranges. We suggest that with moderate *Wolbachia*-mediated blocking and distinct biology, negative-sense viruses represent a fruitful comparator model to other viruses for understanding blocking mechanisms in mosquitoes.

**Author Summary:** Jamestown Canyon Virus (JCV), a newly emerging virus in North America, causes disease when it spills out of its wild mammal hosts into human populations via the bite of infected mosquitoes. We show that the mosquito *Aedes aegypti*, known for transmitting many viral pathogens to humans globally, and whose distribution is creeping northward in the USA toward regions where JCV is present, is likely able to transmit the virus. *Wolbachia* is an endosymbiotic bacterium being released in wild mosquito populations of mosquitoes because it limits the replication of human viruses inside the mosquito, limiting their transmission to humans. We show that *Wolbachia* has a limited ability to control the replication of JCV, which is likely because *Wolbachia*-induced antiviral response is quite weak, and unique aspects of negative-sense virus biology make them less susceptible to blocking. Our findings suggest that JCV may serve as a comparative model to positive-sense viruses like dengue in dissecting the mechanism of *Wolbachia*-mediated virus blocking. It also warns that shifting mosquito distributions, as expected under a changing climate, could bring JCV and Aedes mosquitoes into greater contact, potentially increasing the incidence of JCV in humans.

## Introduction

Jamestown Canyon virus (JCV) is a negative-sense arbovirus in the genus *Orthobunyavirus* (family *Bunyaviridae*), and a relative of La Crosse (LACV) and California Encephalitis viruses. JCV is part of an enzootic cycle involving White-tailed deer, as well as other ungulates, (reviewed in[1]). Transmitted between mammals by mosquitoes, JCV has been found in numerous species including at least 22 members [2] of the genera *Aedes, Culex, Culiseta, Ochlerotatus*, and *Coquillettidia* (reviewed in [3]). Currently, the upper Midwest of the USA, including the states of Minnesota and Wisconsin, has the highest incidence of human JCV (CDC). Seroprevalence in humans in some communities has been found to be as high as 20-40% [2]. Infection is usually asymptomatic but can induce fever, malaise, and headache [1]. Unlike most arboviral illnesses, JCV can also cause respiratory symptoms. In rare instances, the disease becomes neuroinvasive, with death occurring in < 2% of cases [4]. JCV in humans has been on the rise over the last few decades, exhibiting a > 9-fold increase in symptomatic cases. The increase may be explained by better detection and awareness [3,5] as well as factors known to drive enzootic disease spillovers like the destruction of habitat, human encroachment, and climate change [2].

*Aedes aegypti*, and to some extent its sister species *Aedes albopictus*, transmit some of the most prevalent arboviruses globally including dengue (DENV), Zika, Chikungunya, and Yellow Fever [6]. The distribution of both vector species has expanded in recent decades, from tropical to subtropical and even to some temperate regions around the world [7,8]. By 2050, the more tropical *Aedes aegypti* is expected to occur consistently in the upper Midwest [7], where there are already sporadic reports during the summer months (CDC, 2017). In addition, both case and seroprevalence data for JCV in humans and animals suggest the virus is present beyond the upper Midwest, including in the Western, Northeastern, and Southern regions of the USA [3]. To date, JCV has not been isolated from field-caught *Ae. aegypti* or *Ae. albopictus* and there have been no laboratory studies to test for vector competence.

Without deployable vaccines for many of the above viruses, vector control has been the primary means of limiting arboviral diseases in human populations. A recent and promising approach has involved the use of the insect bacterial endosymbiont, *Wolbachia* as a biocontrol agent [9]. *Wolbachia* infection in insects has the effect of limiting the ability of any co-infecting viruses to replicate. This effect combined, with the vertically transmitted microbe’s ability to spread through populations has rendered it a powerful agent against virus transmission.*Wolbachia’s* release into *Ae. aegypti* populations in the field have conferred substantial reductions in dengue fever incidence in human populations living in the release zones in Malaysia, Brazil, and Indonesia [10–12]. Since the discovery of *Wolbachia*-viral mediated blocking, its mechanistic basis has been highly sought after but remains incomplete.

Understanding the basis of the blocking trait is a priority for the effective rollout of this agent across a diverse global landscape and into the future where virus [13] or mosquito [14] evolution may challenge the effectiveness of blocking. The current consensus is that the trait is likely multifaceted (reviewed in [15]), involving diverse aspects of insect physiology, part of which includes the insect immune response, triggered by infection with the bacterial endosymbiont. Specific antiviral pathways and more generalist antioxidant responses [16–18] are part of an innate immune priming response. *Wolbachia’s* ability to block positive-sense RNA viruses including DENV, Zika, chikungunya, and Yellow Fever has been well characterized. In contrast, blocking of the negative-sense viruses LACV, vesicular stomatitis virus (VSV), and cell-fusing agent virus has only been tested *in vitro*, with little evidence of blocking [19,20]. With a poor understanding of whether blocking is mediated locally at the level of the cell, systemically [21,22], or both, vector competence experiments in the whole mosquito are needed for negative-sense viruses. Comparing the virus-blocking ability of *Wolbachia* with different types of viruses may offer a means to further dissect the blocking mechanism.

The innate immune landscape of a mosquito is shaped independently by *Wolbachia* and viral infections, and by their interactions during co-infection. The RNAi and the Toll pathways are considered the most important immune responses against positive-sense DENV in mosquitoes [23,24]. In *Drosophila* RNAi has also been shown to be critical for limiting negative-sense viruses [25]. The antiviral JAK-STAT pathway has also been shown to be critical for controlling DENV [26], and in mosquitoes, it can be activated by Vago protein [27], an interferon-like factor. The activation of Vago through another pathway, IMD, is a likely a strong contributor to the mechanism by which *Wolbachia* reduces loads of co-infecting positive-sense viruses [28,29]. *Wolbachia* infection triggers the expression of both Toll and IMD pathways by inducing reactive oxygen species (ROS) in mosquitoes [16]. Interestingly, in mammals, interferons represent one of the key host defenses against infection with negative RNA viruses, like Rift Valley fever virus [30].

We carried out a vector competence study of JCV in wildtype and *Wolbachia*-infected *Ae. aegypti* to understand the potential emergence of this mosquito species as a vector, and to further explore the limits of *Wolbachia*-mediated blocking. We also examined the innate immune gene expression of *Ae. aegypti* in response to JCV, *Wolbachia*, and co-infection with both, to assess the reaction of the non-native vector to this new virus and the potential role of immunity in blocking negative-sense viruses.

## Results

### Vector competence for Jamestown Canyon Virus

We infected mosquito lines with and without *Wolbachia* infection with Jamestown Canyon Virus (JCV) by oral feeding and then studied viral prevalence and viral load across key tissues (abdomen, head & thorax) at several days post-infection (dpi). The *Wolbachia*-free (wildtype line) showed very strong viral prevalence in mosquito tissues in both replicate experiments, with rates >90% by three dpi (Fig. 1A; S1A Fig), in both the abdomen and head & thorax, indicating dissemination of virus. Bimodal data like these are a common feature of viral loads in mosquitoes [31]. Viral loads averaged across all dpi and both replicate experiments were high; 5.25×10^7^ copies/tissue in abdomen and 8.39 × 10^7^ copies/tissue in the head & thorax (Fig. 1B). Virus prevalence in legs and saliva (Fig. 2A) were near 100% across all dpi, and above 50% after dpi 7, respectively. Saliva estimates of viral prevalence are a conservative estimate of transmissibility [32]. Average viral loads in the legs and saliva (Fig. 2B) were 6.84 ×10^7^ and 1.00 ×10^4^ copies/tissue, respectively. These data strongly demonstrate that, *Ae. aegypti* is highly competent for JCV, allowing rapid virus dissemination through the mosquito body and excretion into the saliva.

**Figure 1.**
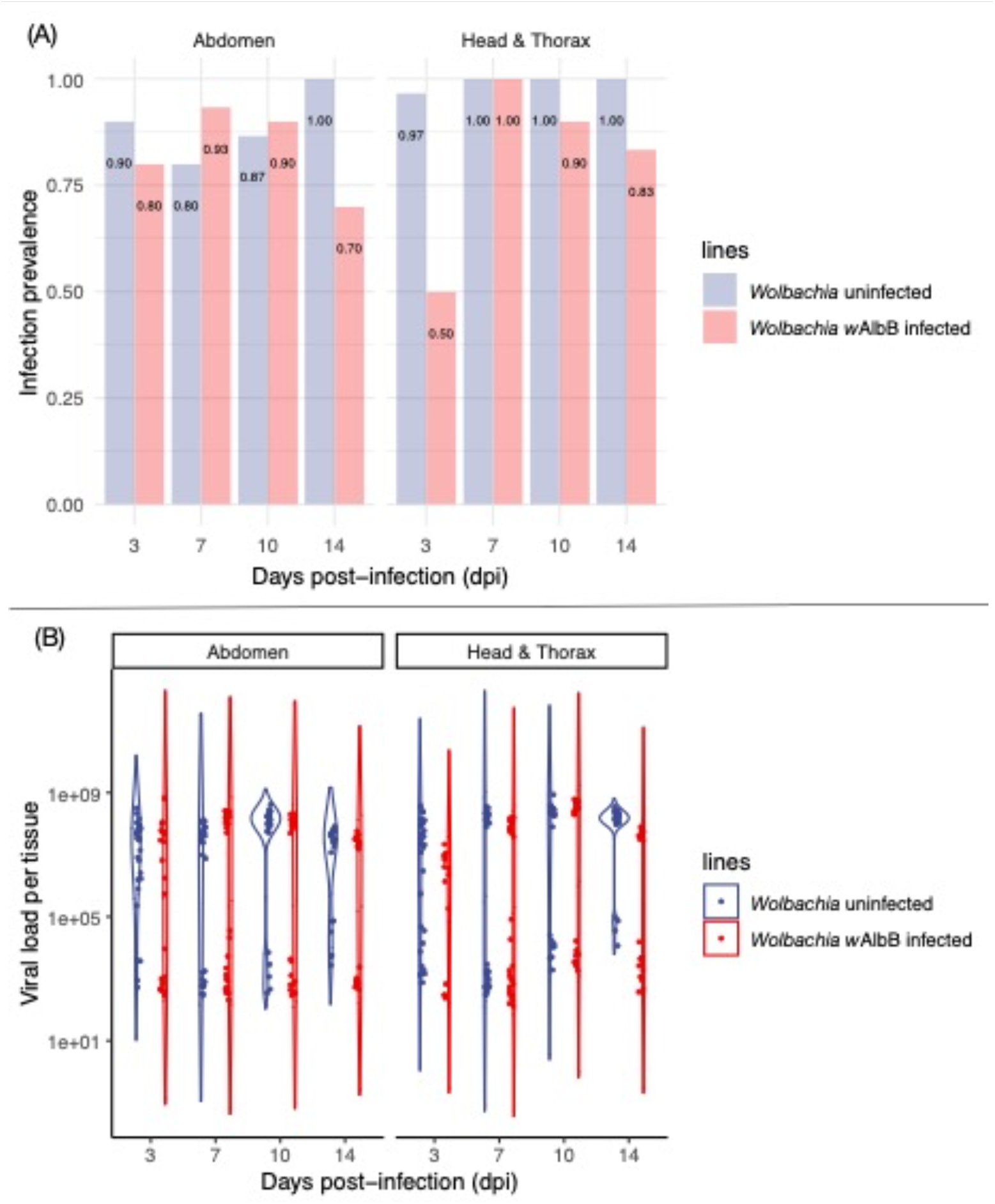
(A) Jamestown Canyon virus (JCV) infection prevalence in adult female mosquito abdomen and head & thorax at 3, 7, 10 and 14 days post-infection (dpi). (B) Quantification of JCV load in adult female mosquito abdomen and head & thorax at 3, 7, 10 and 14 days post-infection (dpi).

**Figure 2.**
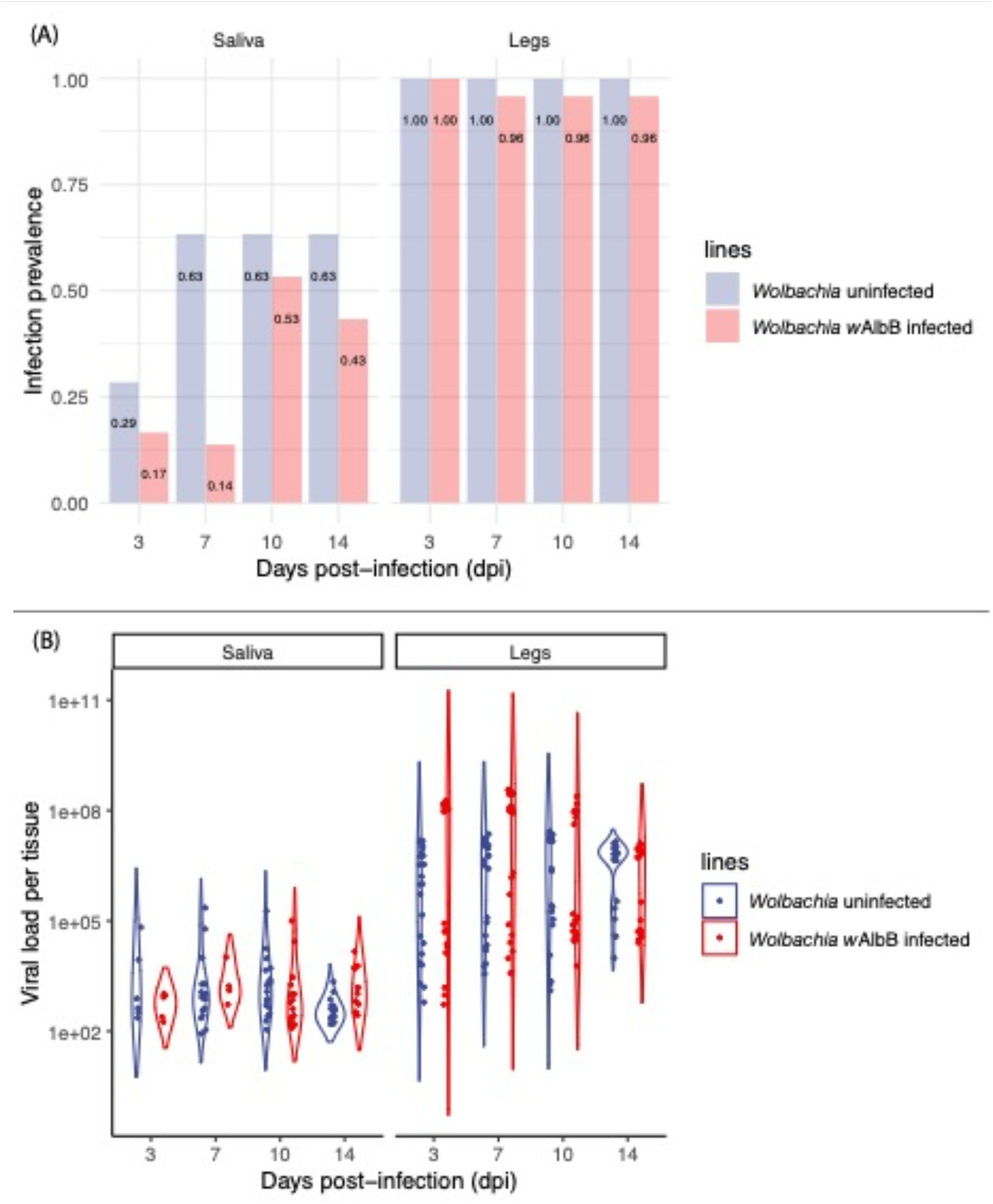
(A) Jamestown Canyon virus (JCV) infection prevalence in adult female mosquito saliva and legs at 3, 7, 10, and 14 days post-infection (dpi). (B) Quantification of JCV load in adult female mosquito abdomen and head & thorax at 3, 7, 10, and 14 days post-infection (dpi).

The effect of *Wolbachia* infection on JCV prevalence (Fig. 1A, S1A Fig.) varied by tissue but was consistent across two replicate experiments (replicates: z = -1.29, p = 0.20; *Wolbachia* infection: z = -5.65, p < 0.001; dpi: z = 5.68, p < 0.001, tissue: z = 1.79, p = 0.074). There was no effect of *Wolbachia* infection in abdomen (z = -1.14, p = 0.25), nor an effect of dpi (z = -0.09, p = 0.93). There was, however, a significant interaction (z = 4.11, p < 0.001), whereby *Wolbachia* reduced prevalence at dpi 3 and 14 in replicate one (Fig. 1A) and dpi 3 (S1A Fig) in replicate two. For head & thorax, *Wolbachia* infection (z = -3.32, p < 0.001), dpi (z = 2.99, p = 0.003), and the interaction (z = 3.19, p = 0.001) were all significant. Like in abdomen, *Wolbachia* infection showed the greatest ability to reduce the prevalence of JCV at 3 and 14 dpi (Fig. 1A) in replicate one and 3 dpi in replicate two (S1A Fig.). When *Wolbachia* infection was associated with reduced prevalence, reductions ranged from 20-50%.

The effect of *Wolbachia* on viral load (Fig. 1B, S1B Fig.) varied across the two replicate experiments (replicate: t = 4.49, p < 0.001; *Wolbachia* infection: t = -0.48, p = 0.63; dpi: t = 1.74, p = 0.082, tissue: t = 3.83, p < 0.001). We, therefore, analyzed the replicates separately. For replicate one (Fig. 1B), *Wolbachia* had no effect in the abdomen (t = -0.68, p = 0.49), nor dpi (t = -0.67, p = 0.51), but had a significant interaction (t = 4.80, p < 0.001). Specifically, *Wolbachia* led to decreased loads in JCV at dpi 14 in replicate 1 and day 3 in replicate 2 (S2 Table). For head & thorax, *Wolbachia* infection (t = -2.34, p = 0.020), dpi (t = 3.04, p = 0.002) and the interaction (t = 2.57, p = 0.011) were all significant. *Wolbachia* infection reduced viral loads on average by a factor of 0.68 on average across all time points in the head & thorax, with the greatest effect at day 3 (S2 Table). In contrast, for replicate two (S1B Fig.), in either tissue *Wolbachia* (abdomen: t = 0.22, p = 0.82; head & thorax: t = 1.94, p = 0.053) or dpi (abdomen: t = 1.84, p = 0.066; head & thorax: t = -0.97, p = 0.33) had no effect on viral load, but a significant interaction was found for head & thorax but not for abdomen (abdomen: t = 0.22, p = 0.83; head & thorax: t = 2.67, p = 0.008). Similar to replicate 1, the greatest reduction in JCV in association with *Wolbachia* infection was at dpi 3 (S2 Table).

Saliva and legs were tested in addition to measure JCV dissemination efficiency, an approximation of viral transmissibility. The prevalence of virus in saliva was affected by *Wolbachia* infection (z = -3.66, p < 0.001) and dpi (z = 3.57, p < 0.001), and with a significant interaction (z = -2.36, p = 0.018). *Wolbachia* infection was more likely to reduce viral load in the early time points (Fig. 2A). In legs there was no effect of *Wolbachia* infection (z = - 0.01, p = 0.995) or dpi (z = - 0.78, p = 0.43). For viral load in saliva (Fig. 2B), there was no effect of dpi (t = -1.03, p = 0.31) or *Wolbachia* infection (t = - 0.69, p = 0.49). Viral load in legs (Fig. 2B) was significant for both dpi (t = - 3.64, p < 0.49) and *Wolbachia* infection (t = 6.07, p < 0.49). On average, across all time points, *Wolbachia* infection reduced infection prevalence by 0.23% in saliva and 3% in legs and reduced viral load by 50% in saliva but unexpectedly increased it by around 9.5 times in legs.

### Effect of Jamestown Canyon Virus infection on immune gene expression

We selected five mosquito immune genes whose expression has been shown to control virus infections and/or be responsive to *Wolbachia* infection. We then tested their relative expression levels in *Wolbachia w*AlbB infected and uninfected lines, with and without JCV infection at different dpi (Fig. 3). Mosquito samples from the same line fed with mock-infected blood served as controls. Although the response of these pathways varied significantly by dpi, our results showed that the RNAi pathway (represented by gene *AGO*) was the major responder to JCV infection, as well as *Vago* (encodes an interferon-like factor), with expression levels increasing ^∼^1000-fold at 10 dpi in both lines. We also found that the JAK/STAT pathway (represented by *hop*) demonstrated a 100-fold increase in expression at 14 dpi in the Wildtype (*Wolbachia*-free) line. AGO, Vago, and hop showed rising trends with increasing dpi (Fig. 3). *MYD88* and *IMD* which represent Toll and IMD pathways, respectively, exhibited very little expression level change (< 10-fold) in response to JCV infection. When each gene was considered separately, we found all genes showed significant differences at different dpi (*MYD88*: F_3,247_ = 41.71, p < 0.001; *IMD*: F_3,247_ = 23.20, p < 0.001; *hop*: F_3,247_ = 9.24, p < 0.001; *AGO*: F_3,247_ = 77.45, p < 0.001; *Vago*: F_3,247_ = 78.57, p < 0.001). Specifically, *hop* (F_3,247_ = 28.24, p = 0.018), *AGO* (F_3,247_ = 4.97, p = 0.027), and *Vago* (F_3,247_ = 16.02, p < 0.001) showed higher expression in the wildtype versus *Wolbachia*-infected lines. *Wolbachia* had no effect on the expression of gene *MYD88* (F_3,247_ = 1.203, p = 0.274) and *IMD* (F_3,247_ = 1.589, p = 0.209) after mosquito infection with JCV. In general, the impact of *Wolbachia* on mosquito immune gene expression was smaller in comparison to JCV (S3 Fig, S4 Table), reducing the expression of the JAK/STAT pathway and increasing the expression of the IMD pathway up to 5-fold dpi.

**Figure 3.**
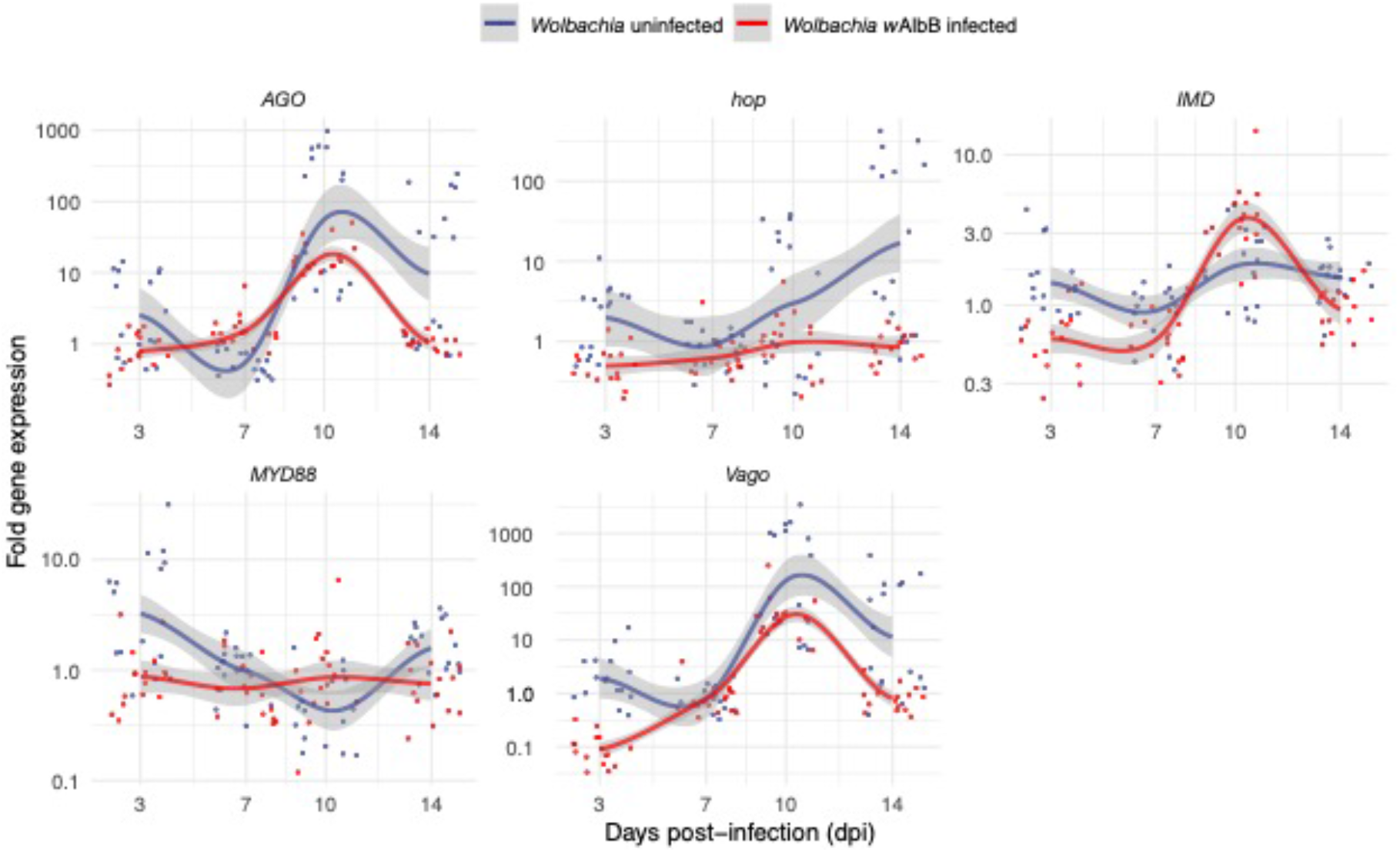
Fold expression change for immunity genes in response to Jamestown Canyon virus infection via 2^−ΔΔCt^ method. Mosquitoes fed with mock-infected blood served as controls. Points are jittered. Smooth curves are plotted using the loess method to indicate the change trend, the gray area indicates 95% interval.

### Effect of Jamestown Canyon virus infection on *Wolbachia* density and female longevity

We quantified the relative density of *Wolbachia* for the whole body of *w*AlbB-infected mosquitoes at different days post-feeding with JCV-infected or mock-infected blood (Fig. 4). We found there was no effect of JCV infection (F_1,120_ = 3.78, p = 0.054), a slight effect of dpi (F_3,120_ = 2.78, p = 0.044), and no interaction (F_3,120_ = 2.35, p = 0.08). *Wolbachia* significantly reduced survival (Fig. 5) by pair-wise log-rank tests whether females had been fed with JCV (χ2 = 14.5, df = 1, p < 0.001) or mock (χ2 = 15.3, df = 1, p < 0.001). JCV infection had no effect on female survival for either *Wolbachia* infected (χ2 = 0.4, df = 1, p = 0.52) and uninfected (χ2 < 0.1, df = 1, p = 0.89) lines.

**Figure 4.**
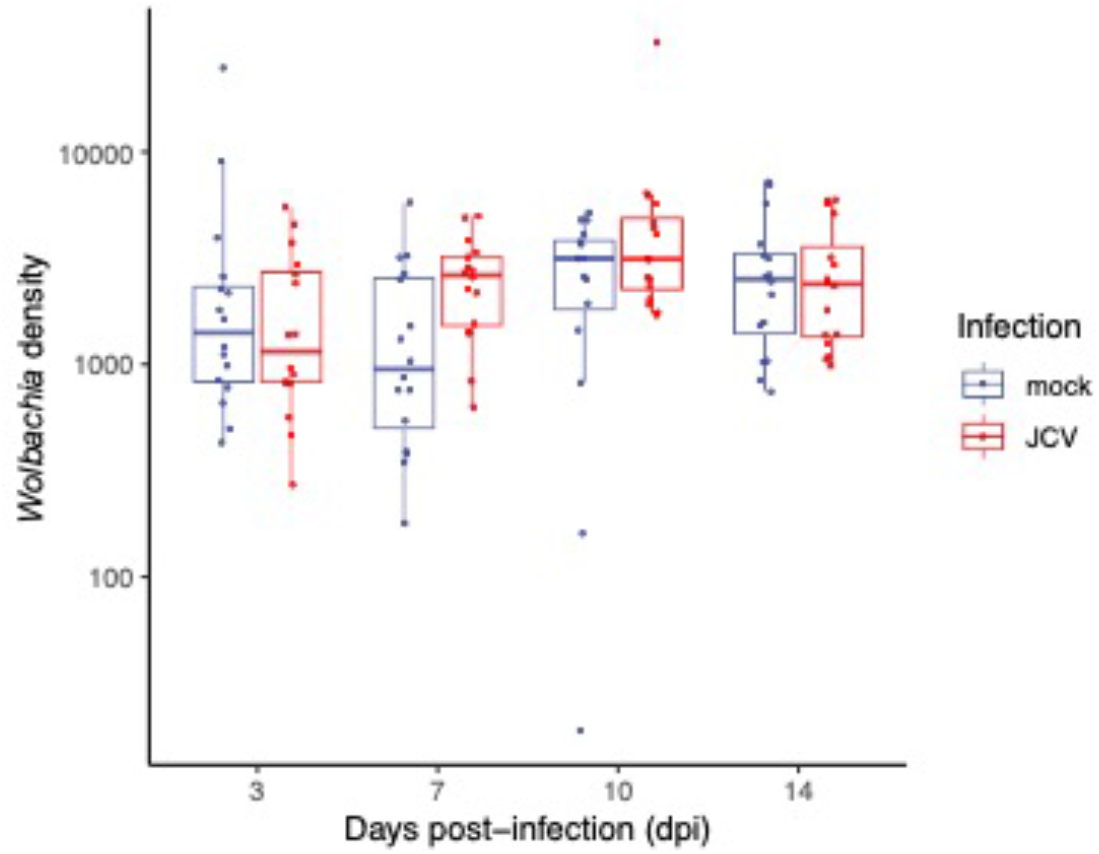
Relative *Wolbachia* density in whole *w*AlbB-infected *Ae. aegypti* at various days post blood feed with either JCV or mock.

**Figure 5.**
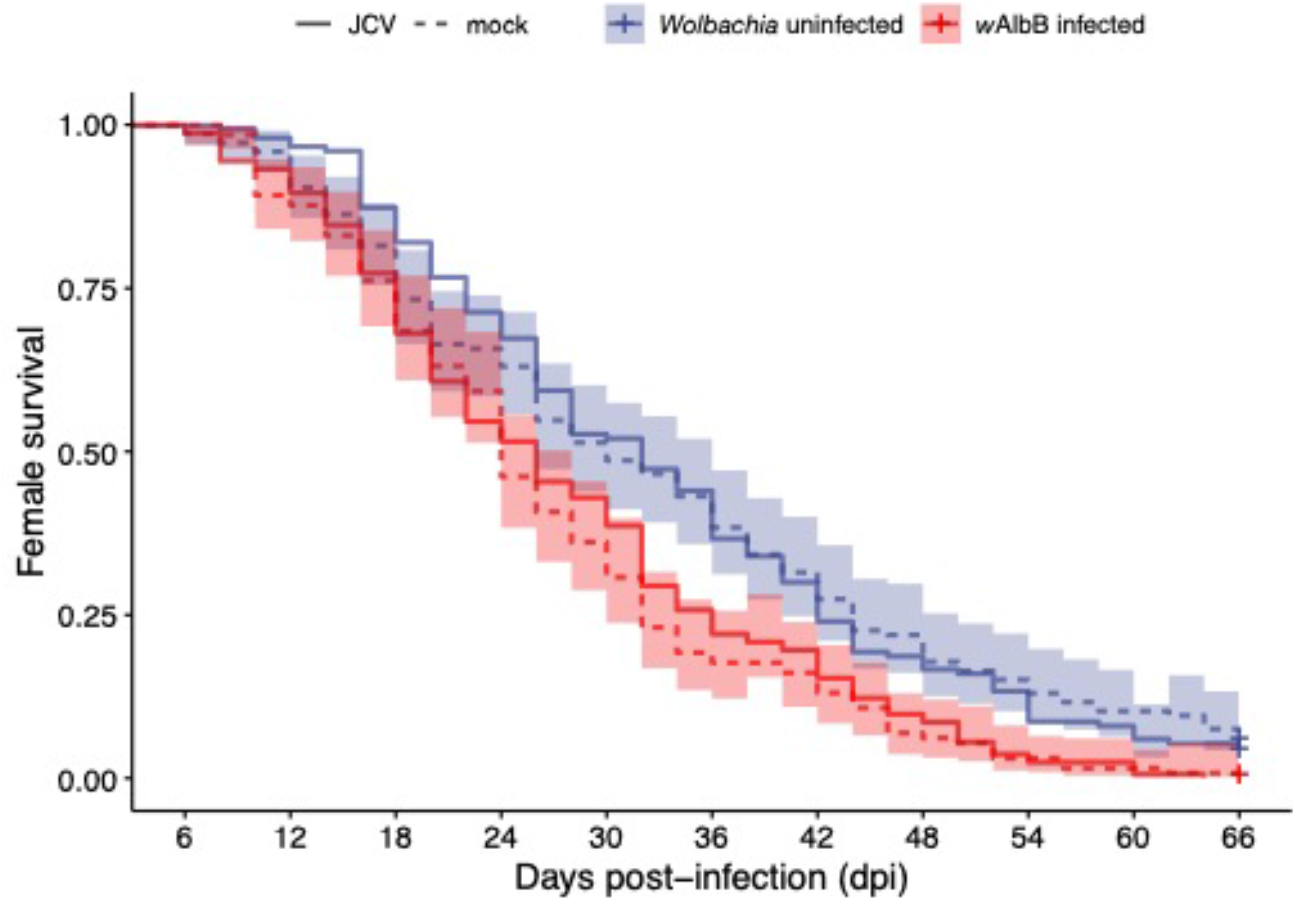
Longevity of *Wolbachia*-infected and uninfected *Ae. aegypti* females post blood feed with either JCV or mock.

## Discussion

In this study, we tested the vector competence of *Ae. aegypti* for Jamestown Canyon Virus (JCV) and investigated whether there is *Wolbachia*-mediated blocking against this negative-sense RNA virus. We demonstrated *Ae. aegypti’s* potential to be a highly competent vector given the following: 1) the infection prevalence and viral loads were very high, near 100% and 10^7^ viral copies/tissue, respectively, and 2), the virus disseminated rapidly, showing dissemination and presence in the saliva at only 3 days post-infectious feed. Additionally, the virus did not reduce female mosquito longevity, leaving this aspect of vectorial capacity unaffected, and lifetime transmission potential high. In terms of the impact of *Wolbachia*, we found that *Wolbachia* only had a moderate ability to inhibit JCV, reducing virus prevalence and viral loads, largely very early during infection.

The emergence and spread of infectious diseases of zoonotic origin have become crucial issues in global human health in recent decades [33,34]. Study shows that JCV, now circulating primarily in the upper Midwest [1,3] has the potential to become a much larger public health threat if its distribution collides with the expanding ranges of *Ae. aegypti* [7] and likely, is close relative, *Ae. albopictus*. Most of the current vectors of JCV are likely feeding on both humans and animals [2], directly facilitating human spillover infections. *Ae. aegypti*, in contrast, feeds with near-perfect fidelity on humans [35]. Its presence in the region would therefore not facilitate spillover but could drive a direct human-human transmission cycle if the virus were sufficiently common. *Ae. albopictus*, in contrast, is more flexible in its host range, depending on availability. It can feed exclusively on humans [35] or other mammals including deer [36] and hence could be a potential participant in both an enzootic and anthroponotic cycle.

Interestingly, *Ae. albopictus* naturally harbors *Wolbachia* infection [37], unlike, *Ae. aegypti* where *Wolbachia* had to be artificially introduced [38], but our work suggests the symbiont may do little to limit the vector competence of wild *Ae. albopictus* for JCV. JCV loads in the body of the mosquito were very high (averaging 10^7^ viral copies/disseminated tissues), following a starting total load of 10^6^ viral copies/midgut. These values are on the higher end of what is seen for similarly designed studies with DENV in *Ae. aegypti*, which is a natural vector (reviewed in [39]). Little is known about the range of JCV titers in the blood of infected humans or wild mammals. For the closely related LACV, titers resulting from artificial infections in deer, and other mammals ranged from 10^2^ to 10^5^/ml [40,41]. Even if titers are low in human and wild hosts, *Ae. aegypti* is very effective at amplifying and disseminating JCV.

*Ae. aegypti* mounted a very strong immune response to JCV that peaked at 10 dpi, but with infection prevalence and viral loads continuing to remain high. This may suggest poor efficacy of the response, or more likely, a steady state between virus replication and anti-viral immune action. In the case of DENV infection where kinetics have been more closely examined, viral loads also plateau and remain high and constant [31] despite a well-described anti-DENV response [42]. The immune response largely consisted of the RNAi and JAK/STAT pathways. While negative-sense viruses do not make dsRNAs in sufficient quantities to trigger RNAi [43], regardless, this pathway has been shown in *Drosophila* to both respond to and limit negative-sense viruses [44]. The parallel expression patterns of *AGO* and *Vago* in response to infection are likely due to molecular crosstalk. *Vago*, produced by the IMD pathway or *Wolbachia* infection, encodes an interferon-like factor that is also a mediator of both the JAK/STAT [29] and RNAi pathways [45]. Last, JCV infection did not appear to shorten mosquito lifespan, suggesting the virus is not cytotoxic in this novel host and that there aren’t extreme fitness tradeoffs for mounting the strong immune response seen here (reviewed in [46]). This is perhaps not surprising as LACV in its native vector *Aedes triseriatus* [47] has no discernable effect on fitness and the virus can live for very long periods in the mosquito, including in the ovaries. The latter is associated with transovarial transmission and overwintering of the virus in northern climates [47].

*Wolbachia*-mediated blocking is known to be a multifaceted blocking trait [15], whose composition may vary depending on the *Wolbachia* strain, virus species/genotype, and vector species/genotype. Previous studies have suggested that viral RNA early in infection is the primary target for blocking [48], possibly recognized by the immune system [16,49], but that *Wolbachia*-induced changes in mosquito physiology involving lipid trafficking [50], autophagy [50,51], and reactive oxygen species [16], etc can also play a role. Dissecting the relative contributions of these multiple components has been challenging in a tripartite organismal system, with no ability to genetically modify *Wolbachia* and difficulties in modifying DENV. The presence of *Wolbachia*-mediated blocking of JCV in both tissues and saliva early in infection offers new comparative potential for dissecting blocking mechanisms relative to positive-sense viruses. The innate immune pathways including RNAi [18], IMD (including *Vago*), and Toll are involved with *Wolbachia*-mediated blocking of positive-sense viruses [17]. Relative to JCV infection, however, *Wolbachia* only moderately induced immune gene expression. Furthermore, during co-infection, *Wolbachia* appeared also to dampen the immune induction by JCV at some time points (see *AGO, vago*, and *hop*, as well as *MYD88* and *IMD* at 3 dpi) suggesting immune activation is unlikely to be at play here.

We propose two hypothetical models for why *Wolbachia*-mediated blocking may be less able to control JCV than positive-sense viruses. First, LACV is known to cause a dramatic shut-off of host protein synthesis in BHK cells, and then subsequently, viral protein synthesis in the later stages of infection [52]. This strategy is thought to protect viruses from attack by host antiviral factors. Mosquito cells infected with LACV continue to divide and behave normally despite infection and so appear unaffected. Interestingly, *Wolbachia* infection has a similar effect on host protein synthesis, reducing it by 23% in *Drosophila* JW18 cells [53], possibly contributing to virus blocking. Negative-sense viruses utilize their own polymerase to produce mRNA copies of their genomes and the activity level of this polymerase is regulated by the amount of viral protein present. If there was some *Wolbachia*-associated inhibition of protein synthesis, therefore, JCV would be expected to overproduce mRNAs in response. Positive-sense viruses, without the mRNA intermediate, would not be affected. Second, other members of the *Orthobunyaviridae*, in mammalian cells, have been shown to remove the 5’ terminus of their genome post-transcriptionally to prevent detection by receptors that trigger interferon activity [54]. There may be parallel stealth activities in the vector to avoid triggering the action of the interferon-like factor Vago [55]. This is difficult to assess, however, with many aspects of the insect antiviral pathways still undefined including key receptors, and modes of action. When examining *Wolbachia* infection alone (S3 Figure), the strongest immune activation was for *Vago* early during infection. The impact of *Wolbachia’s* activation of *Vago* may be greater for DENV and other positive sense viruses where viral infection induction of the gene is much less strong than for JCV [56]. Last, we suggest that studying the *Wolbachia*-mediated blocking seen here early during infection may reveal the functional importance of more universal aspects of blocking not specific to virus type.

In summary, we show that *Ae. aegypti* has the potential to be a very effective vector of JCV in the future, with the mosquito’s range shifting northward [7], and JCV being present beyond the Midwest. Future studies should explore the vector competence of *Ae. albopictus* for JCV, which given its more temperate range, may be a more imminent threat. Additionally, characterizing JCV titers in wild mammals and humans may help to provide context for the vector competence seen here. We also show some evidence of *Wolbachia*-mediated blocking of a negative-sense virus inside the mosquito, whereas previous *in vitro* studies with other viruses found no evidence of blocking [19,20]. This suggests that JCV may serve as a good comparator for teasing out both aspects of blocking that may vary by virus sense-specific biology and indeed those that are more universal. We propose two hypothetical scenarios that may be explored experimentally *in vitro* and *in vivo*. This work should revive interest in studying negative-sense viruses in mosquitoes as a model for *Wolbachia*-mediated blocking.

## Material and methods

### Mosquito rearing

We used two mosquito lines in our experiments. A wildtype, *Wolbachia*-free line (*W*-) derived from a lab-established colony of wildtype mosquitoes collected ^∼^5 years prior in Mérida, Mexico. The second line consisted of a laboratory colony of *Ae. aegypti* infected with the *w*AlbB (*W*+) strain of *Wolbachia* [37]. All mosquitoes were maintained in a climate-controlled insectary, at 26 ± 1°C and 68 ± 5% relative humidity, with a 12:12 hour light : dark cycle. Larvae were reared in plastic trays (40 × 30 × 8cm) containing deionized (DI) water at a density of 300 larvae/3L of water and fed with one tablet of Tropical Fish Food (Tetramin: Tetra Werke, W. Germany) every two days. Adult mosquitoes received 10% sucrose solution *ad libitum*, and females were blood-fed weekly with human blood from anonymous donors (BioIVT: Westbury, NY) using a Hemotek membrane feeder (Hemotek Ltd., UK) when eggs were required for experimental or colony rearing purposes.

### Infection of mosquitoes with Jamestown Canyon virus

Vero cells were cultured at 37 °C (5% CO_2_) with Dulbecco’s Modified Eagle Medium (DMEM) containing 10% FBS (Fetal bovine serum) and 1% Penicillin-Streptomycin (Pen-Strep). The Jamestown Canyon Virus (strain CT 15-8-72591 – passage history = Vero 2) was obtained from the UTMB Arbovirus Reference collection. When cells reached 80% confluency, the media was removed and replaced with new media containing 2% FBS. Cell culture supernatants were collected 3 days post-infection, and the number of virus gene copy numbers was determined through RT-qPCR. Virus-infected cell supernatant was then re-suspended 1:1 in fresh whole human blood and used in an infectious blood meal. 6 ±1 day-old adult mosquitoes were immobilized on ice and females were spread into 2 L paper containers (Zoro, Cat. no. G4215843) and were starved for ^∼^18 hours before feeding. Females were fed on the blood-virus mixtures or blood-mock mixtures (1 : 1) for 40 minutes using the Hemotek membrane feeder system to keep the blood warm at 37°C. After feeding, females were immobilized on ice the second time, and engorged females were sorted into 16 oz paper containers (Zoro, Cat. no. G4141713) and provided with 10% sucrose solution ad libitum through a mesh lid.

At 0 dpi, whole females from both the *Wolbachia* infected or uninfected lines were collected in 2 mL micro tubes containing a ceramic bead and 300 μL of TRI Reagent^®^ (Sigma-Aldrich, Cat. no. T9424). These individuals were then used to assess initial viral load post-feeding (S5 Table), with individuals averaging ^∼^10^6^ viral copies per individual. The remaining fully engorged females were separated into containers for collection at 3, 7, 10, and 14 days post-infection (dpi). In the first replicate, at each time point we collected samples of females from four tissues: wings + legs, saliva, abdomen and head + thorax. The saliva samples were first collected using a previously established capillary method [57]. All samples were stored in 300μL of TRI Reagent^®^, kept at -80 °C until further processing. In the second independent replicate, we only tested abdomen and head + thorax. Infection rates in the abdomen were explored to capture early infection events, including virus in the midgut tissue, whereas head/thorax measures served as proxies for dissemination. Legs and saliva measurements are used to indicate the potential transmissibility of virus and the result from legs might be more accurate [32].

### Jamestown Canyon virus quantification

For nucleic acid extraction, samples were homogenized as previously described [58] using a Bead Ruptor Elite (Omni International, Kennesaw, GA). Trizol (Invitrogen) extraction of total RNA from individual whole mosquitoes was performed following manufacturer’s instructions, resuspended in nuclease-free water, and quantified using the NanoDrop 2000 spectrophotometer system (ThermoFisher Scientific). JCV levels per tissue were quantified by RT-qPCR using a LightCycler^®^ 480 instrument (Roche), the copies of JCV per reaction were multiplied by ten to obtain the copies of JCV per tissue. The primer set was designed by Kinsella et al [4] and quantified by SYBR Green assay (S4 Text).

### *Wolbachia* qualification and gene expression assays

After infectious blood feeding, whole body *Wolbachia* positive and negative females were collected at 3, 7, 10 and 14 dpi, stored in 300μL of TRI Reagent^®^, kept at -80 °C for confirmation of their total body virus loads (S5 Fig.) DNA and RNA were extracted from all samples collected above using a Direct-zol DNA/RNA Miniprep kit (Zymo Research, Cat No. R2080), and RNA was used for gene expression, while DNA was used for *Wolbachia* quantification followed the methods described previously [1]. Relative *Wolbachia* density was based on a 2^-ΔCt^method using mosquito (RPS17 forward: 5’-TCCGTGGTATCTCCATCAAGCT-3’, reverse: 5’-CACTTCCGGCACGTAGTTGTC-3’) and *Wolbachia* (Walb forward: 5’-CCTTACCTCCTGCACAACAA-3’, reverse: 5’-GGATTGTCCAGTGGCCTTA -3’) primer sets that described previously [14]. The q-PCR cycle was activated at 95 °C for 30 seconds (Ramp Rate: 4.4 °C/s), followed by 40 amplification cycles and melting curve analysis same as the viral quantification assays described in S6 Text.

To test the immune response of female *Wolbachia* positive and negative *Ae. aegypti* after Jamestown Canyon virus infection, we selected four genes: *MYD88, IMD, hopscotch* and *AGO2* to represent the immune response from the Toll pathway [16], IMD pathway [16], JAK/STAT pathway [26,59], and RNAi pathway [60] respectively. In addition, we also tested the expression level of *AeVago1*, which encodes Vago protein, an interferon-like factor. The q-PCR was run under the same conditions as above for virus quantification. We used gene *rps17* to normalize expression levels through the 2^-ΔΔCt^ method [61]. Primers used were listed in S7 Table.

**Table 1.** Current evidence from the literature on role of immune genes in response to *Wolbachia* and RNA virus infection.

| Gene ID | pathway | Responds to *Wolbachia*? | Inhibits positive sense RNA viruses? |
| --- | --- | --- | --- |
| *MYD88* (AAEL007768) | Toll pathway | Yes [17] | Yes [62] |
| *IMD* (AAEL010083) | IMD pathway | Yes [17] | unknown |
| *hopscotch* (AAEL012553) | JAK/STAT pathway | unknown | Yes [26] |
| *AGO2* (AAEL017251) | RNAi pathway | unknown | Yes [63] |
| *AeVago1* (AAEL000200) | encode Vago protein | Yes [28] | Yes [28] |
| *rps17* (AAX84650) | 40S ribosomal protein S17 | Reference gene |  |

### Female longevity

The longevity of *Aedes aegypti* females was tested under the infection of *Wolbachia* and/or JCV virus (that fed with JCV or its mock media). After feeding, engorged females were immobilized on the ice and isolated in groups of 14-15 females in 16 oz containers. Each line was represented by eight to eleven cups. Mosquitoes were fed through mesh lids using 2 cotton balls, one soaked in 10% sucrose and another with only water, that were changed every 2 days. Female survival was checked every two days from 4 to 66 days post-infectious feeding.

### Statistical analysis

We used R v. 4.2.1 to conduct data analyses and visualizations. For virus quantification and dissemination, we used a generalized linear model to test the infection prevalence in different tissues and a generalized additive model to test the viral load after JCV infectious blood feeding. The relative gene expression and *Wolbachia* density data were log 10 transformed for ANOVA analysis. Longevity data was compared using the Log-Rank survival test. We also provide pair-wise comparisons (S2 Table and S4 Table) between *Wolbachia* infected and uninfected strains using Mann-Whitney-U-Test (for viral load), Chi-squared test (for infection prevalence) or Tukey’s HSD test (gene expression).

## References

1. Coleman KJ, Chauhan L, Piquet AL, Tyler KL, Pastula DM. An Overview of Jamestown Canyon Virus Disease. Neurohospitalist. 2021;11: 277–278. doi:10.1177/19418744211005948

2. Hollis-Etter KM, Montgomery RA, Etter DR, Anchor CL, Chelsvig JE, Warner RE, et al. Environmental conditions for Jamestown Canyon virus correlated with population-level resource selection by white-tailed deer in a suburban landscape. PLoS One. 2019;14: e0223582. doi:10.1371/JOURNAL.PONE.0223582

3. Pastula DM, Hoang Johnson DK, White JL, Dupuis AP, Fischer M, Staples JE. Jamestown Canyon Virus Disease in the United States-2000-2013. Am J Trop Med Hyg. 2015;93: 384–389. doi:10.4269/AJTMH.15-0196

4. Kinsella CM, Paras ML, Smole S, Mehta S, Ganesh V, Chen LH, et al. Jamestown Canyon virus in Massachusetts: clinical case series and vector screening. Emerg Microbes Infect. 2020;9: 903. doi:10.1080/22221751.2020.1756697

5. Grimstad PR, Shabino CL, Calisher CH, Waldman RJ. A case of encephalitis in a human associated with a serologic rise to Jamestown Canyon virus. Am J Trop Med Hyg. 1982;31: 1238–1244. doi:10.4269/AJTMH.1982.31.1238

6. Campbell LP, Luther C, Moo-Llanes D, Ramsey JM, Danis-Lozano R, Peterson AT. Climate change influences on global distributions of dengue and chikungunya virus vectors. Philos Trans R Soc Lond B Biol Sci. 2015;370: 1–9. doi:10.1098/RSTB.2014.0135

7. Kraemer MUG, Reiner Jr. RC, Brady OJ, Messina JP, Gilbert M, Pigott DM, et al. Past and future spread of the arbovirus vectors Aedes aegypti and Aedes albopictus. Nat Microbiol. 2019/03/06. 2019;4: 854–863. doi:10.1038/s41564-019-0376-y

8. Kraemer MU, Sinka ME, Duda KA, Mylne AQ, Shearer FM, Barker CM, et al. The global distribution of the arbovirus vectors Aedes aegypti and Ae. albopictus. Elife. 2015/07/01. 2015;4: e08347. doi:10.7554/eLife.08347

9. Flores HA, O’Neill SL. Controlling vector-borne diseases by releasing modified mosquitoes. Nat Rev Microbiol. 2018/05/20. 2018;16: 508–518. doi:10.1038/s41579-018-0025-0

10. Nazni WA, Hoffmann AA, NoorAfizah A, Cheong YL, Mancini M V, Golding N, et al. Establishment of Wolbachia Strain wAlbB in Malaysian Populations of Aedes aegypti for Dengue Control. Curr Biol. 2019/11/26. 2019;29: 4241–4248 e5. doi:10.1016/j.cub.2019.11.007

11. Gesto JSM, Pinto SB, Dias FBS, Peixoto J, Costa G, Kutcher S, et al. Large-Scale Deployment and Establishment of Wolbachia Into the Aedes aegypti Population in Rio de Janeiro, Brazil. Front Microbiol. 2021/08/17. 2021;12: 711107. doi:10.3389/fmicb.2021.711107

12. Utarini A, Indriani C, Ahmad RA, Tantowijoyo W, Arguni E, Ansari MR, et al. Efficacy of Wolbachia-Infected Mosquito Deployments for the Control of Dengue. N Engl J Med. 2021/06/10. 2021;384: 2177–2186. doi:10.1056/NEJMoa2030243

13. Edenborough KM, Flores HA, Simmons CP, Fraser JE. Using Wolbachia to Eliminate Dengue: Will the Virus Fight Back? J Virol. 2021;95. doi:10.1128/JVI.02203-20

14. Ford SA, Allen SL, Ohm JR, Sigle LT, Sebastian A, Albert I, et al. Selection on Aedes aegypti alters Wolbachia-mediated dengue virus blocking and fitness. Nat Microbiol. 2019/08/28. 2019;4: 1832–1839. doi:10.1038/s41564-019-0533-3

15. Lindsey ARI, Bhattacharya T, Newton ILG, Hardy RW. Conflict in the Intracellular Lives of Endosymbionts and Viruses: A Mechanistic Look at Wolbachia-Mediated Pathogen-blocking. Viruses. 20180321st ed. 2018;10. doi:10.3390/v10040141

16. Pan X, Zhou G, Wu J, Bian G, Lu P, Raikhel AS, et al. Wolbachia induces reactive oxygen species (ROS)-dependent activation of the Toll pathway to control dengue virus in the mosquito Aedes aegypti. Proc Natl Acad Sci U S A. 2011/11/30. 2012;109: E23–31. doi:10.1073/pnas.1116932108

17. Pan X, Pike A, Joshi D, Bian G, McFadden MJ, Lu P, et al. The bacterium Wolbachia exploits host innate immunity to establish a symbiotic relationship with the dengue vector mosquito Aedes aegypti. ISME J. 2018;12: 277–288. doi:10.1038/ISMEJ.2017.174

18. Terradas G, Joubert DA, McGraw EA. The RNAi pathway plays a small part in Wolbachia-mediated blocking of dengue virus in mosquito cells. Sci Rep. 2017/03/07. 2017;7: 43847. doi:10.1038/srep43847

19. Schultz MJ, Tan AL, Gray CN, Isern S, Michael SF, Frydman HM, et al. Wolbachia wStri Blocks Zika Virus Growth at Two Independent Stages of Viral Replication. mBio. 20180522nd ed. 2018;9. doi:10.1128/mBio.00738-18

20. Schnettler E, Sreenu VB, Mottram T, McFarlane M. Wolbachia restricts insect-specific flavivirus infection in Aedes aegypti cells. J Gen Virol. 2016;97: 3024–3029. doi:10.1099/JGV.0.000617

21. Osborne SE, Iturbe-Ormaetxe I, Brownlie JC, O’Neill SL, Johnson KN. Antiviral protection and the importance of Wolbachia density and tissue tropism in Drosophila simulans. Appl Environ Microbiol. 2012;78: 6922–6929. doi:10.1128/AEM.01727-12

22. Amuzu HE, McGraw EA. Wolbachia-Based Dengue Virus Inhibition Is Not Tissue-Specific in Aedes aegypti. PLoS Negl Trop Dis. 20161117th ed. 2016;10: e0005145. doi:10.1371/journal.pntd.0005145

23. Franz AWE, Sanchez-Vargas I, Adelman ZN, Blair CD, Beaty BJ, James AA, et al. Engineering RNA interference-based resistance to dengue virus type 2 in genetically modified Aedes aegypti. Proc Natl Acad Sci U S A. 2006;103: 4198–4203. doi:10.1073/PNAS.0600479103

24. Ramirez JL, Dimopoulos G. The Toll immune signaling pathway control conserved anti-dengue defenses across diverse Ae. aegypti strains and against multiple dengue virus serotypes. Dev Comp Immunol. 2010;34: 625–629. doi:10.1016/j.dci.2010.01.006

25. Muellera S, Gausson V, Vodovar N, Deddouchea S, Troxler L, Perot J, et al. RNAi-mediated immunity provides strong protection against the negative-strand RNA vesicular stomatitis virus in Drosophila. Proc Natl Acad Sci U S A. 2010;107: 19390–19395. doi:10.1073/PNAS.1014378107

26. Souza-Neto JA, Sim S, Dimopoulos G. An evolutionary conserved function of the JAK-STAT pathway in anti-dengue defense. Proc Natl Acad Sci U S A. 2009;106: 17841–17846. doi:10.1073/PNAS.0905006106

27. Paradkar PN, Duchemin JB, Voysey R, Walker PJ. Dicer-2-dependent activation of Culex Vago occurs via the TRAF-Rel2 signaling pathway. PLoS Negl Trop Dis. 2014;8. doi:10.1371/JOURNAL.PNTD.0002823

28. Asad S, Parry R, Asgari S. Upregulation of Aedes aegypti Vago1 by Wolbachia and its effect on dengue virus replication. Insect Biochem Mol Biol. 2018;92: 45–52. doi:10.1016/J.IBMB.2017.11.008

29. Mukherjee D, Das S, Begum F, Mal S, Ray U. The Mosquito Immune System and the Life of Dengue Virus: What We Know and Do Not Know. Pathogens. 2019;8. doi:10.3390/PATHOGENS8020077

30. Ly HJ, Ikegami T. Rift Valley fever virus NSs protein functions and the similarity to other bunyavirus NSs proteins. Virol J. 2016;13. doi:10.1186/S12985-016-0573-8

31. Novelo M, Hall MD, Pak D, Young PR, Holmes EC, McGraw EA. Intra-host growth kinetics of dengue virus in the mosquito Aedes aegypti. PLoS Pathog. 2019/12/04. 2019;15: e1008218. doi:10.1371/journal.ppat.1008218

32. Gloria-Soria A, Brackney DE, Armstrong PM. Saliva collection via capillary method may underestimate arboviral transmission by mosquitoes. Parasit Vectors. 2022;15. doi:10.1186/S13071-022-05198-7

33. Braack L, Gouveia De Almeida AP, Cornel AJ, Swanepoel R, De Jager C. Mosquito-borne arboviruses of African origin: review of key viruses and vectors. Parasit Vectors. 2018;11. doi:10.1186/S13071-017-2559-9

34. Tajudeen YA, Oladunjoye IO, Mustapha MO, Mustapha ST, Ajide-Bamigboye NT. Tackling the global health threat of arboviruses: An appraisal of the three holistic approaches to health. Health Promot Perspect. 2021;11: 371–381. doi:10.34172/HPP.2021.48

35. Ponlawat A, Harrington LC. Blood Feeding Patterns of Aedes aegypti and Aedes albopictus in Thailand. J Med Entomol. 2005;42: 844–849. Available: https://academic.oup.com/jme/article/42/5/844/863877

36. Richards SL, Ponnusamy L, Unnasch TR, Hassan HK, Apperson CS. Host-Feeding Patterns of Aedes albopictus (Diptera: Culicidae) in Relation to Availability of Human and Domestic Animals in Suburban Landscapes of Central North Carolina NIH Public Access. J Med Entomol. 2006;43: 543–551.

37. Xi Z, Khoo CC, Dobson SL. Wolbachia establishment and invasion in an Aedes aegypti laboratory population. Science (1979). 2005/10/15. 2005;310: 326–328. doi:10.1126/science.1117607

38. Walker T, Johnson PH, Moreira LA, Iturbe-Ormaetxe I, Frentiu FD, McMeniman CJ, et al. The wMel Wolbachia strain blocks dengue and invades caged Aedes aegypti populations. Nature. 2011/08/26. 2011;476: 450–453. doi:10.1038/nature10355

39. Souza-Neto JA, Powell JR, Bonizzoni M. Aedes aegypti vector competence studies: A review. Infection, Genetics and Evolution. 2019;67: 191–209. doi:10.1016/J.MEEGID.2018.11.009

40. Amundson TE, Yuill TM, DeFoliart GR. Experimental La Crosse Virus Infection of Red Fox (Vulpes fulva), Raccoon (Procyon Lotor), Opossum (Didelphis Virginiana), and Woodchuck (Marmota Monax). Am J Trop Med Hyg. 1985;34: 586–595. doi:10.4269/AJTMH.1985.34.586

41. Osorio JE, Godsey MS, Defoliart GR, Yuill TM. La Crosse Viremias in White-Tailed Deer and Chipmunks Exposed by Injection or Mosquito Bite. Am J Trop Med Hyg. 1996;54: 338–342. doi:10.4269/AJTMH.1996.54.338

42. Xi Z, Ramirez JL, Dimopoulos G. The Aedes aegypti Toll Pathway Controls Dengue Virus Infection. PLoS Pathog. 2008;4: e1000098. doi:10.1371/journal.ppat.1000098

43. Weber F, Wagner V, Rasmussen SB, Hartmann R, Paludan SR. Double-Stranded RNA Is Produced by Positive-Strand RNA Viruses and DNA Viruses but Not in Detectable Amounts by Negative-Strand RNA Viruses. J Virol. 2006;80: 5059. doi:10.1128/JVI.80.10.5059-5064.2006

44. Cogni R, Ding SD, Pimentel AC, Day JP, Jiggins FM. Wolbachia reduces virus infection in a natural population of Drosophila. Commun Biol. 2021;4. doi:10.1038/S42003-021-02838-Z

45. Paradkar PN, Trinidad L, Voysey R, Duchemin JB, Walker PJ. Secreted Vago restricts West Nile virus infection in Culex mosquito cells by activating the Jak-STAT pathway. Proc Natl Acad Sci U S A. 2012;109: 18915–18920. doi:10.1073/PNAS.1205231109/-/DCSUPPLEMENTAL

46. Lazzaro BP, Tate AT. Balancing sensitivity, risk, and immunopathology in immune regulation. Curr Opin Insect Sci. 2022;50: 100874. doi:10.1016/J.COIS.2022.100874

47. Borucki MK, Kempf BJ, Blitvich BJ, Blair CD, Beaty BJ. La Crosse virus: Replication in vertebrate and invertebrate hosts. Microbes Infect. 2002;4: 341–350. doi:10.1016/S1286-4579(02)01547-2

48. Bhattacharya T, Newton ILG, Hardy RW. Viral RNA is a target for Wolbachia-mediated pathogen blocking. PLoS Pathog. 2020;16. doi:10.1371/JOURNAL.PPAT.1008513

49. Terradas G, Allen SL, Chenoweth SF, McGraw EA. Family level variation in Wolbachia-mediated dengue virus blocking in Aedes aegypti. Parasit Vectors. 2017/12/29. 2017;10: 622. doi:10.1186/s13071-017-2589-3

50. Voronin D, Cook DAN, Steven A, Taylor MJ. Autophagy regulates Wolbachia populations across diverse symbiotic associations. Proc Natl Acad Sci U S A. 2012;109. doi:10.1073/PNAS.1203519109

51. Brackneyid DE, Correa MA, Cozens DW. The impact of autophagy on arbovirus infection of mosquito cells. 2020 [cited 23 Apr 2023]. doi:10.1371/journal.pntd.0007754

52. Raju R, Kolakofsky D. La Crosse virus infection of mammalian cells induces mRNA instability. J Virol. 1988;62: 27–32. doi:10.1128/JVI.62.1.27-32.1988

53. Grobler Y, Yun CY, Kahler Id DJ, Bergman Id CM, Id HL, Oliver B, et al. Whole genome screen reveals a novel relationship between Wolbachia levels and Drosophila host translation. [cited 24 Apr 2023]. doi:10.1371/journal.ppat.1007445

54. Habjan M, Andersson I, Klingströ J, Schü Mann M, Martin A, Zimmermann P, et al. Processing of Genome 59 Termini as a Strategy of Negative-Strand RNA Viruses to Avoid RIG-I-Dependent Interferon Induction. 2008 [cited 24 Apr 2023]. doi:10.1371/journal.pone.0002032

55. Cheng G, Liu Y, Wang P, Xiao X. Mosquito Defense Strategies against Viral Infection. Trends Parasitol. 2016;32: 177–186. doi:10.1016/j.pt.2015.09.009

56. Asad S, Parry R, Asgari S. Upregulation of Aedes aegypti Vago1 by Wolbachia and its effect on dengue virus replication. 2017 [cited 26 Apr 2023]. doi:10.1016/j.ibmb.2017.11.008

57. Dutra HLC, Rocha MN, Dias FBS, Mansur SB, Caragata EP, Moreira LA. Wolbachia Blocks Currently Circulating Zika Virus Isolates in Brazilian Aedes aegypti Mosquitoes. Cell Host Microbe. 2016;19: 771–774. doi:10.1016/J.CHOM.2016.04.021

58. Dutra HLC, Ford SA, Allen SL, Bordenstein SR, Chenoweth SF, Bordenstein SR, et al. The impact of artificial selection for Wolbachia-mediated dengue virus blocking on phage WO. PLoS Negl Trop Dis. 2021;15. doi:10.1371/JOURNAL.PNTD.0009637

59. Jupatanakul N, Sim S, Angleró-Rodríguez YI, Souza-Neto J, Das S, Poti KE, et al. Engineered Aedes aegypti JAK/STAT Pathway-Mediated Immunity to Dengue Virus. PLoS Negl Trop Dis. 2017;11. doi:10.1371/JOURNAL.PNTD.0005187

60. Blair CD. Mosquito RNAi is the major innate immune pathway controlling arbovirus infection and transmission. Future Microbiol. 2011;6: 265–277. doi:10.2217/fmb.11.11

61. Ginzinger DG. Gene quantification using real-time quantitative PCR: An emerging technology hits the mainstream. Exp Hematol. 2002;30: 503–512. doi:10.1016/S0301-472X(02)00806-8

62. Ramirez JL, Dimopoulos G. The Toll immune signaling pathway control conserved anti-dengue defenses across diverse Ae. aegypti strains and against multiple dengue virus serotypes. Dev Comp Immunol. 2010;34: 625–629. doi:10.1016/J.DCI.2010.01.006

63. Sánchez-Vargas I, Scott JC, Poole-Smith BK, Franz AWE, Barbosa-Solomieu V, Wilusz J, et al. Dengue virus type 2 infections of Aedes aegypti are modulated by the mosquito’s RNA interference pathway. PLoS Pathog. 2009;5. doi:10.1371/JOURNAL.PPAT.1000299

